# Inferring hidden causal relations between pathway members using reduced Google matrix of directed biological networks

**DOI:** 10.1101/096362

**Authors:** José Lages, Dima L. Shepelyansky, Andrei Zinovyev

## Abstract

Signaling pathways represent parts of the global biological network which connects them into a seamless whole through complex direct and indirect (hidden) crosstalk whose structure can change during development or in pathological conditions. We suggest a novel methodology, called Googlomics, for the structural analysis of directed biological networks using spectral analysis of their Google matrices, using parallels with quantum scattering theory, developed for nuclear and mesoscopic physics and quantum chaos. We introduce the reduced Google matrix method for the regulatory biological networks and demonstrate how its computation allows inferring hidden causal relations between the members of a signaling pathway or a functionally related group of genes. We investigate how the structure of hidden causal relations can be reprogrammed as the result of changes in the transcriptional network layer during cancerogenesis. The suggested Googlomics approach rigorously characterizes complex systemic changes in the wiring of large causal biological networks.

## 1 Introduction

The network biology point of view on signaling pathway as a part of complex integrated molecular machinery consists in considering it as a subnetwork embedded into a global molecular network. As a consequence, all properties of the pathway functioning depend on the network context to which it remains connected. Considering only the set of direct causal relations between pathway members (as is frequently the case) neglects the indirect effect of the global biological network changes which may significantly re-wire the pathway topology by introducing implicit (hidden) causal relations. Characterizing such influence of the global network structure on the local network properties and dynamics remains one of the major challenges of systems and network biology [3]. A number of empirical and pragmatic approaches have been suggested recently to address this question [18,28,35,14,19].

Reconstructions of the global directed causal signaling network structure have appeared only recently in the form of comprehensive molecular interaction databases such as SIGNOR[45], SignaLink [24], where the pair-wise relations between molecules are oriented. Large-scale cell type-specific reconstructions of transcriptional causal networks have become possible thanks to appearance of Chip-Seq technology [25,27] or advances in computational methodology of transcription factor binding site predictions combined with the data on chromatin accessibility [41].

In the previous works, quantification of indirect interactions (sometimes called influences) between pathway members mainly exploited the calculation of shortest or second shortest paths (the paths that become shortest after removal of an edge in a shortest path), following quantification of the balance between negative and positive path signs [19,9]. The limitation of such approaches is, however, in that they do not take into account the complex global structure of the network: multiple dense causal connections between two nodes might be more important than a single shortest path connecting them, representing a hypothetical sequence of intermediate regulations.

Global changes in the structure of the global network might effectively re-wire a signaling pathway even if its direct interactions are weakly dependent on the biological context. For example, the functioning of a signaling pathway must be heavily affected by the structure of transcriptional feedbacks indirectly re-wiring the pathway structure by gross effect of implicit (hidden) causal relations.Therefore, changing the transcriptional layer in the global network effectively rewires many signaling pathways even without affecting the structure of direct connections between its members. Therefore, it would be advantageous to develop an efficient and rigorous mathematical formalism allowing quantifying such a phenomenon. In this paper, we suggest a candidate methodology for this purpose.

We assess the characteristics of signal propagation through a pathway by considering the stochastic Markov process of random walk with uniform non-zero restart (teleportation) probability along oriented edges of the graph representing the global biological network. This process is described by Google matrix (see Materials and Methods section), and its stationary state defines PageRank centrality measure of the graph nodes. More complete and subtle description of the process can be obtained by looking at the complete (complex) spectrum of the Google matrix, which might reflect complex non-stationary properties of the random walk. For example, grouped eigenvalues of the Google matrix in the complex plane can define weak communities in the graph where the signaling flow (random walk) can be “trapped” for a finite time. In order to quantify indirect hidden causal relations between the members of a pathway, we introduce the original formalism of reduced Google matrix, based on decomposing the global Google matrix into the parts describing the pathway itself and the influence of the rest of the network.

It is important to note that the Google matrix approach or related random walk-based approaches is now a well established efficient tool for analysis of various complex directed networks including the World Wide Web [10], citation network of Physical Review [43], rating of global importance of scientific journals [6], world trade commercial networks and many other networks [7,20].

To our knowledge, the Google matrix approach or related ideas have been applied before in systems biology only to undirected networks, in order to find activated network modules or to “smooth” the high-throughput data [46,31,34], establish connection of genes to diseases [48,33], improve interpretability of genome-wide analyses [42,38,36] and compute network-based cancer biomarkers [49]. The formalism of reduced Google matrix is applied for biological networks in this paper for the first time.

We describe the details of the Google matrix and reduced Google matrix methodology in the “Methods” section, and in the “Results” section application of the methodology to several large regulatory networks is documented. Using the suggested approach, we quantify the effect of the changes in the structure of transcriptional network as a result of oncogenic events during chronic myelogenous leukemia onto re-wiring connections between proteins in several cancer-related groups of genes. We conclude that the method is able to infer the missing indirect causal relations between the members of a pathway and can detect events of hidden re-wiring during cancerogenesis.

## 2 Results

### 2.1 Used networks and case study description

In order to illustrate the application of Google matrix approach to studying oncogenic changes in the global and local network structures, we constructed two large directed networks describing global signaling in a leukemia cancer cell line K562 compared to a healthy cell line GM12878 derived from normal B-lymphocytes. The transcriptional networks of these two cell lines have been previously characterized [27], using systematic Chip-Seq experiments on a number of transcription factors whose activity is detected in a given cell line. Transcriptional networks for GM12878 and K562 cells have been previously analyzed in order to estimate their structural properties which can lead to buffering and robustness [2]. It was demonstrated that the wiring of the transcriptional network in cancer leads to significant changes in the number of structural patterns leading to violating the network robustness properties. In order to deal with combined signaling+transcriptional networks, we merged each of the transcriptional network to the global reconstruction of signaling taken from the SIGNOR database [45] (version from February 2016). Therefore, as a modeling assumption, we assume that the structure of the global signaling network does not depend on the biological context while the transcriptional regulation layer undergo significant changes leading to indirect effect on the signaling (Figure 1).

**Fig. 1.**
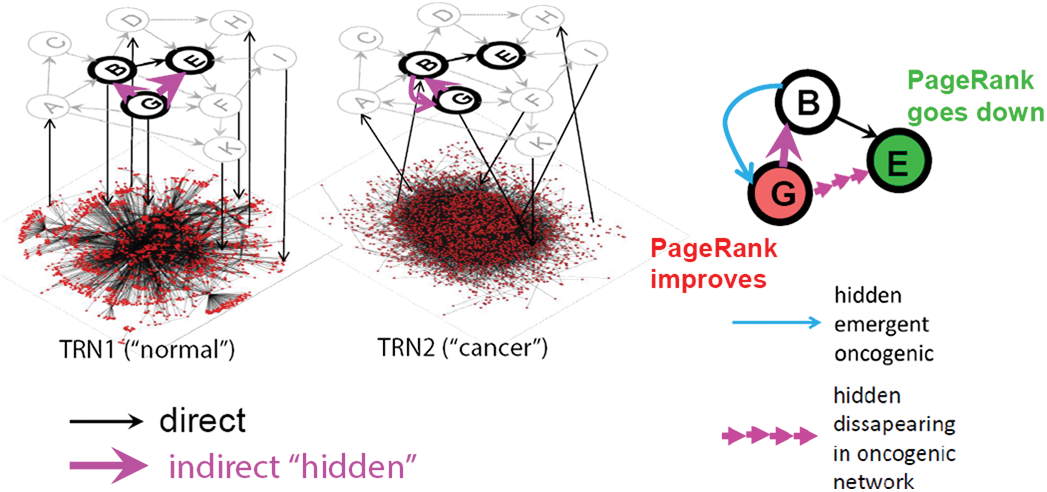
Using reduced Google matrix approach for inferring hidden causal relations in signaling pathways. Here the structure of the context-dependent global regulatory network is symbolically shown as consisting of two layers: the upper (nodes A-K) is the global signaling network whose structure does not depend on the context and the lower is a symbolic view of the contextual transcriptional regulatory network (TRN) whose structure can change between a “normal” and a “cancer” cell. Thick node borders denote a pathway embedded into the global signaling network. Black arrows denote direct physical interactions. Pink arrows denote inferred hidden directed regulations through the global regulatory network (both layers). In the final representation of the pathway (on the right), one can show those hidden regulations which emerge or disappear due to the changes in the TRN structure. Also, the color of the pathway nodes can show the direction of PageRank change: green corresponds to the PageRank decreased in the cancer network while red corresponds to the opposite.

### 2.2 Biological interpretation of PageRank and CheiRank centrality measures and their changes in cancer

#### 2.2.1 Distribution of proteins on PageRank vs CheiRank plane

For the three directed biological networks described above (SIGNOR alone and two merged signaling+transcriptional regulatory networks), we applied the Google matrix methodology as described in Materials and Methods section, and determined the values of PageRank and CheiRank for all proteins. The distribution of proteins in PageRank vs CheiRank plane is shown in Figure 2. This figure shows that in the case of these networks PageRank and CheiRank measures are not correlated which reflects quite distinct biological role of proteins with many incoming and with many outgoing directed interactions.

**Fig. 2.**
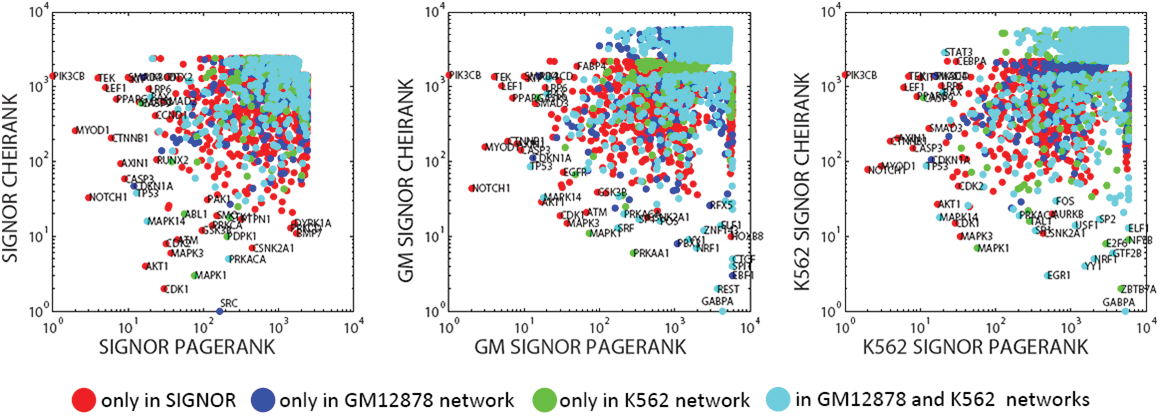
Distribution of proteins on PageRank x CheiRank plane for three networks (SIGNOR - signaling network, GM network is a merge of SIGNOR and GM12878 normal blood cell transcriptional network, K562 network is a merge of SIGNOR and K562 leukemia cancer cell transcriptional network). The colors signifies to which network a protein originally belongs. Red color signifies proteins which are present only in SIGNOR network and not present in any of the two transcriptional networks. Dark blue color signifies proteins which are present in GM12878 transcriptional network and not present in K562 transcriptional network. Green color signifies proteins which are present in K562 transcriptional network and not present in GM12878 transcriptional network. Cyan color signifies proteins which are present in both K562 and GM12878 transcriptional networks.

It can be easily demonstrated (data not shown) that most of the proteins simultaneously having high values of PageRank and CheiRank (such as AKT1, NOTCH1, CTNNB1, TP53, CDKN1A, ATM, MAPK3, CDK1, EGFR) play an important role in cancer biology. This is, however, a rather trivial observation having in mind that it is known that hubs of the protein interaction networks frequently correspond to cancer-related genes [32,3].

One can also observe that adding a transcriptional network to a signaling network (SIGNOR) significantly changes the top ranked proteins for CheiRank but not for PageRank. This is consistent with the fact that the transcriptional networks are characterized by fan-like structures in which a transcription factor can regulate many (hundreds) of proteins, while the cases when a protein is regulated by so many upstream regulator are relatively rare.

Overall, the general shape of the distribution of proteins in the “normal” GM (Figure 2,middle) and “cancer” K562 (Figure 2,right) networks is similar; nevertheless, there are differences. For example, it can be seen that the top Chei-ranked proteins are not the same in these two networks. There is a region in the PageRank vs CheiRank plane for GM network (Figure 2,middle) occupied by proteins which are present in SIGNOR but not in GM12878 transcriptional network (cluster of green color points for CheiRank and PageRank around 1000). Vice versa, there is a region in K562 network (Figure 2,right) occupied by proteins which are present in SIGNOR but not in K562 transcriptional network (cluster of dark blue points). This observation underlines the fact that both the composition and the wiring topology of “normal” and “cancer” networks have important differences.

#### 2.2.2 Detecting “creative protein elements” by comparing PageRank and CheiRank to simple connectivity

A general statement on PageRank and CheiRank is that they are correlated to in-degree and out-degree of a node [23]. However, this correlation is not perfect as it can be seen in Figure 3. Some proteins significantly deviate from the general dependence trend, so it is interesting to consider what kind of non-local network topologies give unexpectedly high PageRank or CheiRank values despite relatively low connectivity degree. This deviation can be scored by a simple product of the rank and the corresponding degree, i.e., *CE*_*in*_ = *K* ·(*InDegree*(1), *CE*_*out*_ = *K*^*∗*^ · (*OutDegree* + 1).

**Fig. 3.**
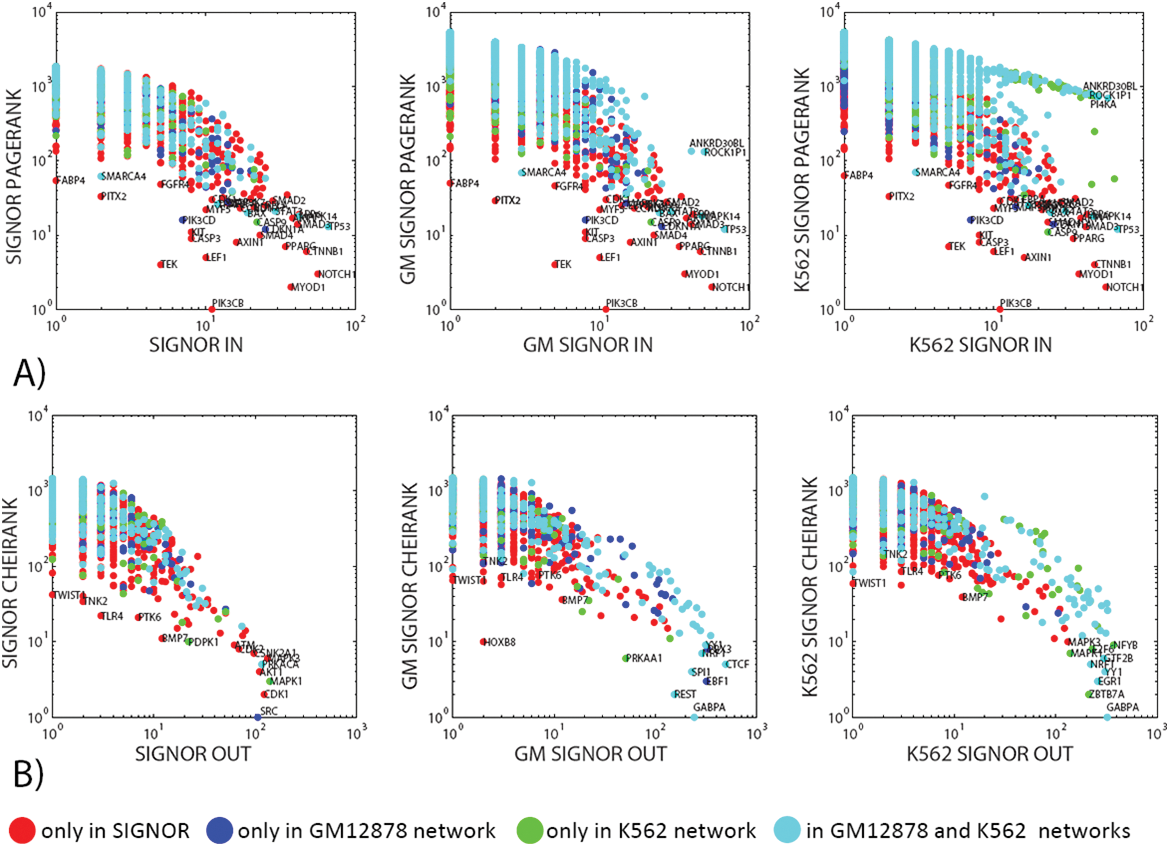
Dependence of PageRank (A) and CheiRank (B) on the in-degree and out-degree correspondingly, in three networks The coloring of nodes corresponds to the description provided in the caption of the Figure 2.

The notion of “creative elements” in the context of biological networks is discussed in [18] as such proteins that are not hubs of the networks themselves but can provide important (and, frequently, transient) connections between network hubs. Also, sometimes, together with hubs, proteins playing the role of “connectors” are discussed as those proteins having high centrality but not connectivity measures [3].

We suggest that the proteins significantly deviating from the the general trend between a Google matrix-based rank and the corresponding connectivity degree are potential candidates for the role of “creative elements” and connectors in the signaling and transcriptional networks. Several examples of such proteins and the local topologies explaining the deviation from the trend are provided in Figure 4.

**Fig. 4.**
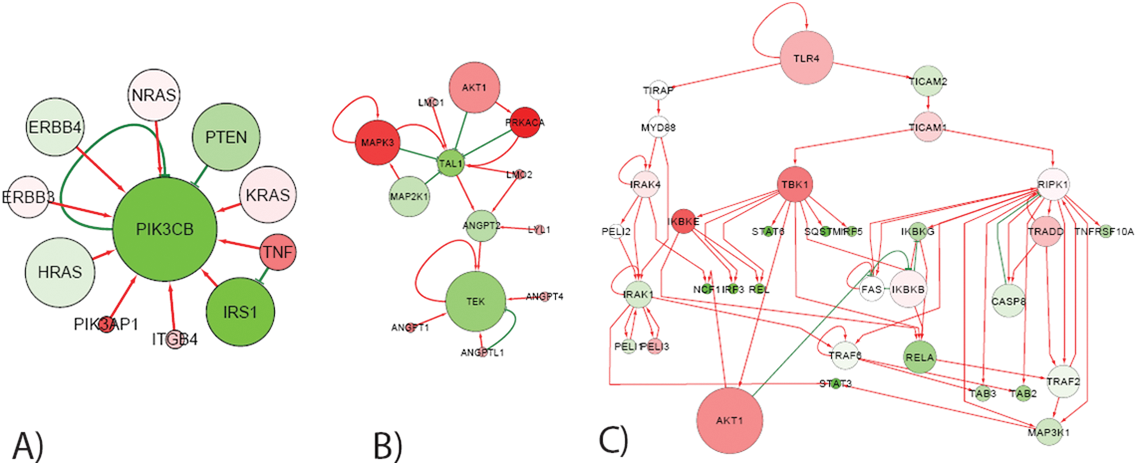
Several examples of network topologies, illustrating emergence of “creative elements” (defined here as proteins whose PageRank or CheiRank can not be simply explained by the number of incoming or outgoing connections correspondingly). A and B) The size of the node is proportional to its PageRank value. C) The size of the protein is proportional to its CheiRank value. The color of proteins in all three panels reflects the ratio between the number of incoming and outgoing edges. If the color is greenish then the protein is “importer” receiving more incoming regulations than having out-going edges. The reddish proteins are “exporters” regulating more proteins downstream than the number of incoming regulations.

For example, the top ranked in PageRank protein in all three networks is PIK3CB (look at Figure 2) having relatively low in-degree (10), which makes it to significantly deviate from the general trend (Figure 4A). PIK3CB gene encodes a catalytic subunit of the kinase PI3K, a key kinase involved in multiple cell signaling cascades. PIK3CB gene is frequently mutated or amplified in several cancer types (such as lung squamous cell carcinoma where its rate of mutations can be as high as 18%), which causes abnormalities in cell survival signaling. From the network point of view, it’s high value of PageRank is probably explained by the fact that, accordingly to SIGNOR, PIK3CB is regulated by several highly connected proteins which have predominantly incoming edges (PTEN, ERBB3, ERBB4, HRAS, NRAS, KRAS, IRS1).

An other example of such a protein is TEK which is ranked #4 by PageRank in SIGNOR network, having only 5 incoming edges. From Figure 4B one can see that TEK is the sink of the cascade TAL→ANGPT2→TEK, which progressively collects incoming regulations, starting from the top connected hubs such as AKT1, MAPK3, PRKACA. The biological function of the TEK protein is quite unique: this is a receptor tyrosine kinase which has several immunoglobulin-like domains, three epidermal growth factor domains and three fibronectin type III repeats in the extracellular part. This makes this protein potential regulator of multiple cellular functions such as angiogenesis, endothelial cell survival, proliferation, migration, adhesion and cell spreading, reorganization of the actin cytoskeleton, and also maintenance of vascular quiescence. Such rich functional cross-talk allows suggesting TEK as a creative element in the global cell signaling network.

Our final example of deviation from the major CheiRank vs out-degree trend is TLR4 protein, which is ranked #22 by CheiRank in SIGNOR having only 2 out-going regulation and 1 self-interaction. From the Google matrix-based network analysis this can be explained by the fact that TLR4 triggers several cascades affecting downstream several major regulators having a large number of out-going edges such as AKT1. Indeed, the biological function of TLR4 (toll-like receptor 4) is activating the innate immune system which requires triggering many important cellular cascades (such as NFkB signaling), regulating a large number of cellular processes.

Other observations from Figure 3 underlies some particular features and differences between “normal” and “cancer” networks. For example, one can notice in Figure 4A,right that in the cancer network there is a large number of proteins with high number of incoming transcriptional regulations but not ranked well by PageRank. This feature is almost absent in the normal GM-SIGNOR network (Figure 4A,middle).

#### 2.2.3 Biological meaning of PageRank and CheiRank changes in cancer

Having seen the differences in the distribution of PageRank and CheiRank in Figure 2 between the “normal” and “cancer” regulatory network, we characterized the relative change of the ranks by computing their log ratio between two networks. Overall, the relative changes in CheiRank had larger amplitude than in PageRank which can be partially explained by the different number of transcriptional targets of the transcriptional factors, described in the GM12878 and K562 transcriptional networks.

We characterized the biological functions represented by those proteins significantly deviating from zero in Figure 5, by applying the standard enrichment analysis based on hypergeometric test, calculating what is the probability (p-value) of a selected protein set to intersect with some predefined protein set by random chance. For this purpose we used the *toppgene* bioinformatics package [13]. For the enrichment analysis, we took those proteins deviating from zero by two standard deviations of the distributions of PageRank of CheiRank log ratios, and analyzed the positive and negative sides of the distribution separately. The results of the analysis are presented in Table 1 and online at http://www.ihes.fr/~zinovyev/googlomics/grn2016/rankchanges/.

**Fig. 5.**
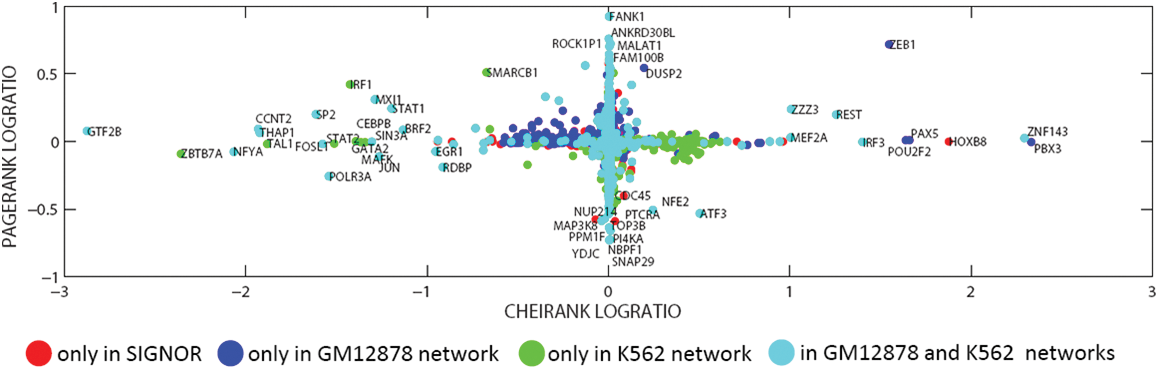
Relative changes in the PageRank and CheiRank in cancer vs normal network. The abscissa is the decimal logarithm of the ratio *K*_*K*562_/*K*_*GM*_, where the subscript denotes in which network the CheiRank is calculated. The ordinate is the decimal logarithm of the ratio 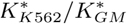. For example, the proteins in the right part of the plot such as MEF2A, ZNF143, ZEB1 are those whose CheiRank significantly increased (i.e., the protein has less outgoing connections) in the “cancer network” K562 (which can be simply because the corresponding transcription factor is not present in K562 transcriptional network, as in the case of ZEB1, blue points in the plot).

**Table 1.**
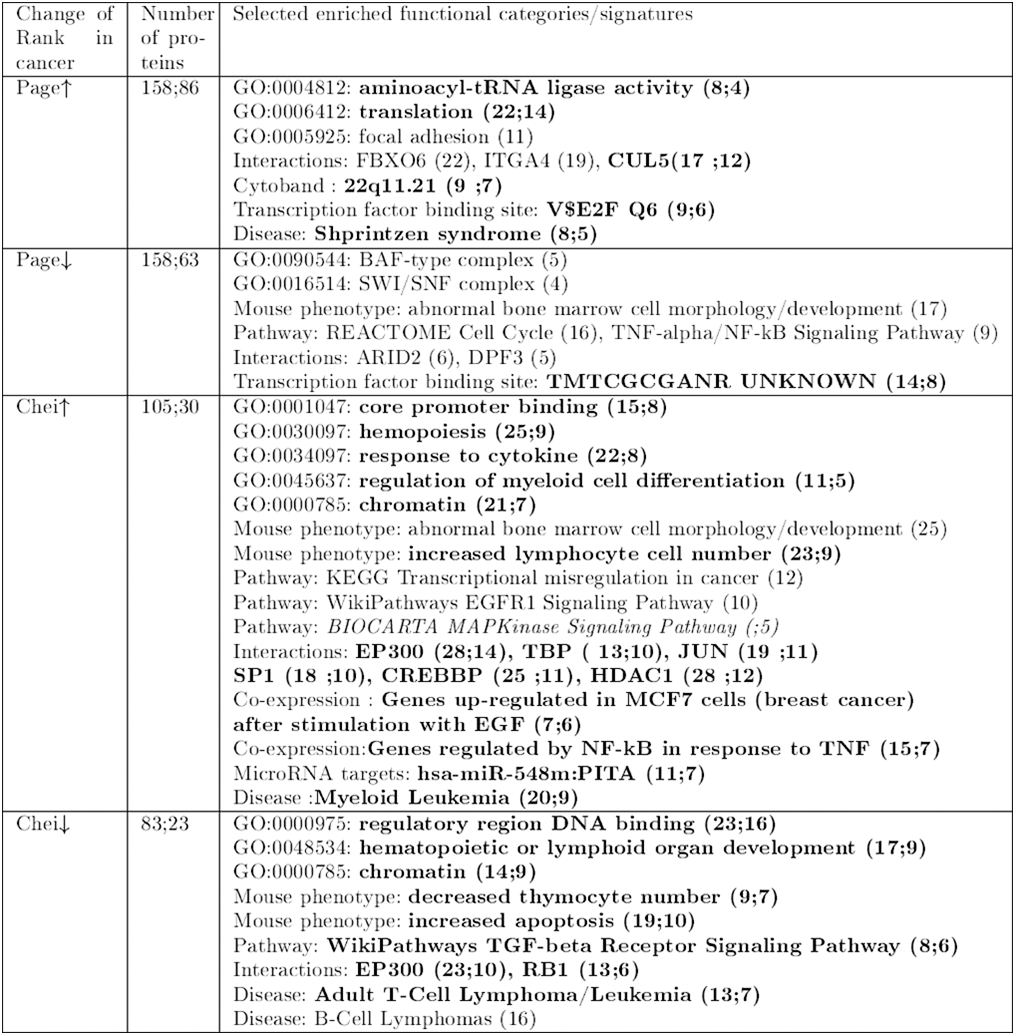
Table of enriched biological functional categories of proteins. The numbers signifies the number of proteins whose ranks are significantly changed. By bold those functional categories are highlighted which found enriched after filtering out proteins not present in either cancer or normal transcriptional networks. In this case, two values for the number of proteins are shown separated by semicolon: one without filtering and one after filtering. Complete interactive version of this table is available at http://www.ihes.fr/~zinovyev/googlomics/grn2016/rankchanges/. Arrow up means that the corresponding rank decreases (or, in other words, “improves”, rank value increases) in the cancer network (the proteins are more connected). Arrow down means the opposite: the corresponding rank “degrades” and the proteins become less connected in cancer.

We’ve noticed, however, that the results of this analysis can be biased by a simple fact that a protein can be included in the “normal” transcriptional network GM12878 (thus having, for example, many transcriptional out-going interactions) and not at all included in the “cancer” transcriptional network K562 (thus having not at all transcriptional out-going interactions). This is the case, for example, for ZEB1 transcriptional factor. Despite the fact that such difference can be “real”, i.e. the transcriptional factor might be not expressed in the case of cancer, hence, does not regulate any genes, we’ve decided to perform an additional analysis focusing only at those proteins which simultaneously present in both “normal” and “cancer” transcriptional networks. These proteins might be present or not in the SIGNOR network. This second analysis made several biological functions detected in the previous analysis insignificant (normal text lines in the Table 1) but many remained significant (bold text lines in the Table 1). For the second analysis, in case of CheiRank, we took those proteins which deviated from zero by one standard deviation in the distribution of log ratio of the CheiRanks, in order to collect a sufficient number of proteins.

Overall, the undertaken analysis shows a picture consistent with the nature of the studied cells. I.e., we show that changes in the CheiRanks between normal and cancer cells highlights a number of proteins previously described as being implicated in leukemia (16 from 53 selected for the second analysis). Interestingly, this analysis highlights proteins implicated in the regulation of myeloid cell differentiation and hemopoiesis, which is also expected. 9 from 30 selected proteins were previously associated with the mouse phenotype characterized by the increased number of lymphocytes and significantly improved their CheiRank in the cancer network (they became more powerfull regulators). Having many transcriptional factors in the transcriptional networks explains significance of such Gene Ontologies as “core promoter binding” and “chromatin” in the analysis of CheiRank changes in both directions, and also interactions with key transcription co-regulators as EP300, HDAC1 and CREBBP.

At the same time, the analysis gives also some unexpected findings. For example, a number of proteins involved in translation (14 from 86 selected) or having the E2F transcription factor binding motif the promoter sequence (6 from 86) or being located in a specific genomic locus 22q11 (7 from 86) improved their PageRank in cancer network (meaning they became more regulated). 8 from 63 genes with a specific motif in their promoter sequences (TMTCGCGANR) showed significant increase in the PageRank (meaning they became less regulated in cancer) is also an unexpected finding.

### 2.3 Inferring hidden causal relations between members of protein sets

Application of Google matrix to the global network allows quantifying the global ranking of protein nodes and their changes, as it was illustrated in the previous section. Reduced Google matrix (see Materials and Methods for formal description) allows focusing on a subset of nodes, quantify local importance of nodes in this subset and also detect indirect (hidden) connections between the members of the subset.

In order to test this approach in the context of biological networks, we’ve defined several protein sets, each of which contains a functionally related group of proteins. However, the meaning of the functional proximity is different in all cases.

We start with a definition of a biological pathway, which play one of the most central roles in all cancer types: AKT-mTOR pathway. It is a molecular cascade downstream of PI3K kinase which is important in regulating the cell cycle and cell survival, controlling a number of normal physiological processes, and being dysregulated in many diseases including cancer. We take the definition of the AKT-mTOR pathway from the external pathway database Atlas of Cancer Signaling Network (ACSN) [37], based on manual mining of molecular biology publications, so the definition of this subset of proteins can be called “knowledge-driven”.

Second analyzed subset is by contrast purely “data-driven” and corresponds to a particular gene expression signature (set of genes), shown to be connected to cell proliferation in multiple cancer studies through data analysis. We used a particular definition of this signature coming from a large multi-cancer study of tumoral transcriptomes, using Independent Component Analysis (ICA) method [51,8]. Gene expression signatures obtained through statistical data analysis sometimes serve as a scaffold for reconstructing the topology of regulatory connections between the corresponding proteins. Therefore, we considered this set for determining direct and indirect connections between its members.

Third analyzed group of proteins is a set of known direct targets of a transcriptional factor E2F1 which is central to regulation and progression through the cell cycle. The member names of this set were manually extracted from reading the molecular biology literature on the functioning of cell cycle in order to reconstruct it as a biochemical reaction diagram [11]. In this case, the challenge is to understand what biological pathways can be directly regulated by E2F1, what are the possible direct and indirect feedbacks to the regulation of E2F1 itself and how they are changing in cancer progression.

All three subsets are central to the studied in this paper cancer progression and its influence on the structure of biological networks. In all three cases, we roughly equilibrated the sizes of the protein sets, limiting them to approximately 50 proteins.

It happened that the direct interactions between the members of all three sets of proteins are described in SIGNOR pathway database, and not in the transcriptional networks. Therefore, one might consider that the wiring of direct connections between the set members are not affected by the changes in the transcriptional program. However, we further show that the structure of indirect connections might change accordingly to the changes in the global context created by the transcriptional network.

#### 2.3.1 AKT-MTOR pathway

From SIGNOR database, we’ve retrieved 138 direct regulatory connections between 63 proteins of AKT-mTOR pathway. These direct connections formed a large connected subnetwork containing the majority (43 proteins) of the pathway members, and the rest was orphan nodes not connected to any other.

No direct transcriptional regulatory connections was found between the members of the pathway; hence, the structure of direct connections did not change in the “cancer” network with respect to the “normal” network.

We’ve computed indirect regulatory relations using the reduced Google matrix approach as described in Materials and Methods section, separately for SIGNOR, the “normal” and “cancer" global regulatory networks, combined the common signaling SIGNOR part and the specific transcriptional network. The strength of the indirect regulation can be evaluated by looking at its *G*_qr_ value.

For the “normal” network we found that the distribution of the corresponding 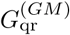 values contains essentially close to zero values, with only some pointing to existence of indirect regulation. Thus, for an arbitrarily chosen threshold *G*_qr_ > 0.01 one detects 50 indirect interactions, ten top of them are shown in Figure 6,A (in magenta color). These 50 indirect regulations connect 8 more proteins into the large connected component.

**Fig. 6.**
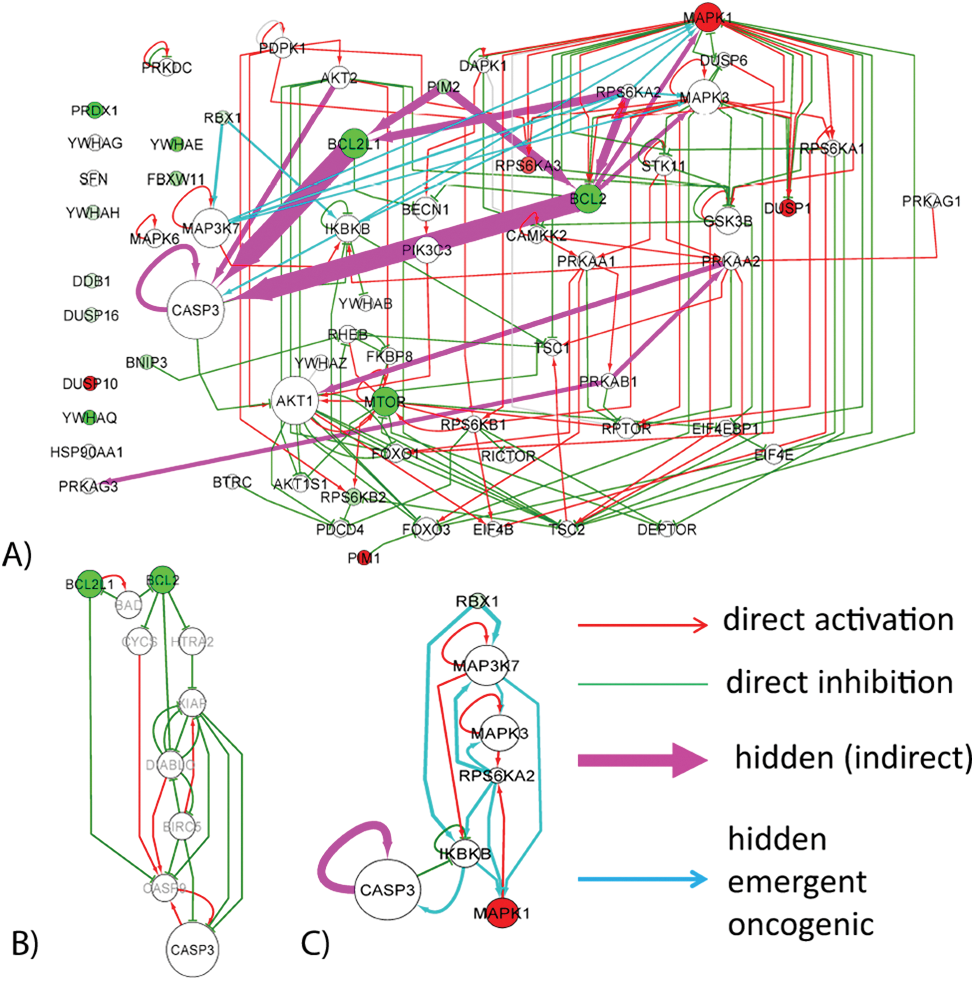
AKT-mTOR pathway reconstructed using SIGNOR database and by inferring indirect connections using reduced Google matrix approach. A) Direct connections representing activator and inhibitor regulations are shown as red and green arrows correspondingly. Magenta color arrows shows the inferred indirect interactions which are common between the “normal” and “cancer” networks. Light blue color arrows represent those indirect interactions which are inferred only in the “cancer” network. Line width of the arrows representing the indirect interactions, is proportional to their *G*qr score. Here only 10 top scored indirect connections present in both “normal" and “cancer" networks and 10 top “emergent in cancer" connections are shown. The color of protein nodes reflects the relative change in their local PageRanks in “cancer" vs “normal" networks. Red color means that the protein become better ranked in PageRank in the “cancer" network, and the green color means the opposite. The size of the node is proportional to the value of the PageRank in the SIGNOR network. B) Hidden cascade of indirect regulations connecting BCL2 and BCL2L1 proteins with CASP3. The proteins shown with grey labels are those which are not present in the definition of AKT-mTOR pathway. C) Cascade of hidden interactions emerging in the “cancer" network and not present in “normal" network.

It can be noticed that the pattern of the top indirect connections is highly non-random and forms two “hidden patwhays”, one pointing to CASP3 protein through BCL proteins, and one connecting PRKA proteins to AKT1. Both hidden pathways have rather clear biological interpretation. The hidden regulations also point out to the important crosstalk between BCL2 and MAPK1, MAPK3 proteins, not represented by direct interactions inside AKT-mTOR pathway.

The first one can be related to the existence of apoptotic pathway in the global regulatory network, where BCL2 and BCL2L1 proteins play an important role. CASP3 serves the final point of the apoptotic pathway, being the main executor protein of the apoptotic process (the executor caspases take care of destroying the proteins of the suicided cell and the cell itself). In order to illustrate how BCL proteins and CASP3 are connected through the global network, we’ve computed the shortest and the second shortest oriented path between BLC2 and BCL2L1 proteins and CASP3 protein (Figure 6,B). It can be seen that these paths include the main players of the apoptotic machinery (XIAP,CASP9,DIABLO,CYCS). Second hidden pathway connects the subunits of AMP-activated protein kinase (AMPK), an important energy sensor protein, to AKT1. As a conclusion, one can state that the reduced Google matrix approach was able to point out to biologically important and meaningful indirect connections between several AKT-mTOR pathway members.

In addition, we compared the inferred indirect interactions in “normal” and “cancer” global networks. We found that there is a strong correlation between all three set of values 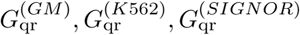 (correlation coefficients are close to 0.998). All strong indirect interactions inferred using the “normal” GM network were also found in “cancer” K562 network. However, in the “cancer” network we found additional candidates for indirect interactions, top ten of which are shown in Figure 6,A and Figure 6,C. It can be seen that such “emergent oncogenic” indirect interactions also underline existence of a “hidden” causal relation between RBX1 and MAPK1 proteins.

#### 2.3.2 Data-driven signature of proliferation-related proteins

We analyzed a set of 49 proteins found in SIGNOR database whose expression was shown to significantly change between fast proliferative and slow proliferative tumors in 9 cancer types [8]. We found 47 direct interactions connecting them into one large connected component consisting of 31 proteins who were predominantly the phosphorylation targets of the cyclin-dependent kinase CDK1, so the structure of the network of direct interactions has a starlike structure organized around one large hub protein. As in the previous example, no direct transcriptional connections were found between the members of this protein set.

We’ve computed indirect regulatory relations using the reduced Google matrix approach as described in Materials and Methods section, separately for SIGNOR, the “normal” and “cancer” global regulatory networks, combined the common signaling SIGNOR part and the specific transcriptional network. As before, we’ve found only a minor fraction of all pair-wise protein relations as candidates for indirect interactions (only 32 from 2305 passed the threshold *G*_qr_ > 0.01). In Figure 7 we show all indirect regulations inferred in the “normal" GM network. As before, with indirect connections, it was possible to connect more proteins (43 out of 49). Thus, the most important indirect connection connects PCNA protein to the largest connected component of the network. PCNA (proliferating cell nuclear antigen) protein is a key cell cycle protein important both for DNA replication and DNA repair.

**Fig. 7.**
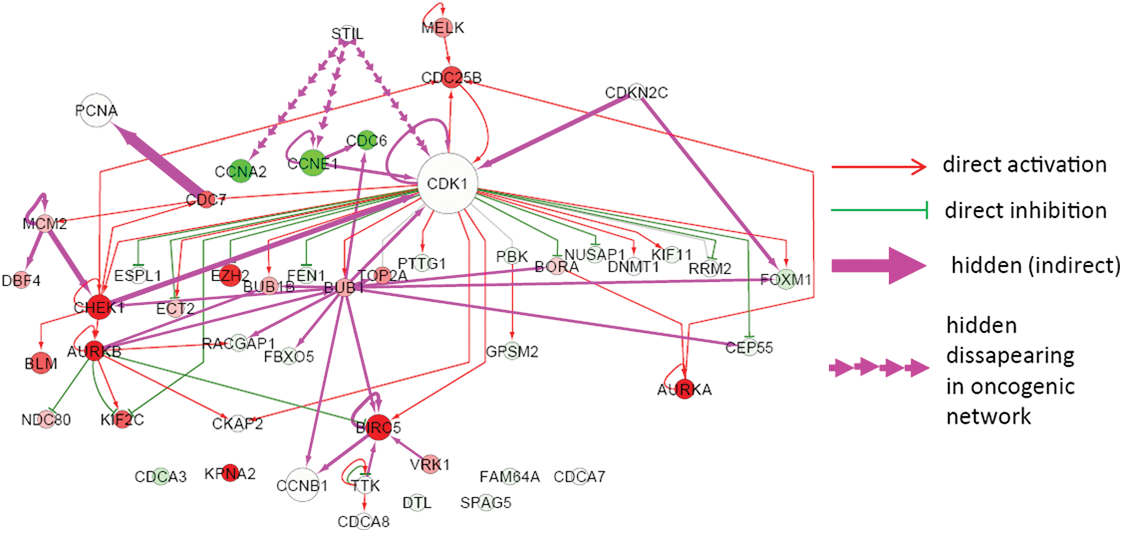
Network of proteins shown to be related to cell proliferation by transcriptomic data analysis. The meaning of the node and edge colors is the same as in Figure 6 besides those regulations which disappear in “cancer" network compared to “normal" network (shown by interrupted line arrows).

While comparing “cancer" and “normal" networks, unlike the previous example, we do not find new “emergent oncogenic” indirect interactions. Instead, we observed that several indirect interactions disappear in the “cancer" network, namely three indirect regulations connecting STIL protein to CCNA2, CCNE1 and CDK1. STIL is a cytoplasmic protein implicated in regulation of the mitotic spindle checkpoint, a regulatory pathway that monitors chromosome segregation during cell division to ensure the proper distribution of chromosomes to daughter cells. Interestingly, STIL protein was shown to be heavily deregulated in T-cell leukemias through genome modifications leading to gene fusions. Disappearance of indirect connections between STIL and CDK1 can be interpreted as loosening the control over several important cyclins (CCNA2 and CCNE1) and the key cell cycle protein CDK1 in cancer. Consistently, we find that the PageRank of the aforementioned cyclins increases (e.g., for the local subnetwork PageRanks, 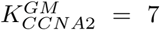 and 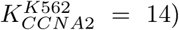 which means that they are less regulated/controlled in cancer. Several other proteins such as AURKA, AURKB, CHEK1, BIRC5, CDC25B decreases their local PageRanks in cancer which means they become more controlled. Interestingly, one of the direct targets of CDK1 kinase, protein BUB1, becomes a new hub of the indirect interactions. Indeed, this protein plays a central role in mitosis by phosphorylating members of the mitotic checkpoint complex and activating the spindle checkpoint.

#### 2.3.3 Set of transcriptional targets of E2F1 transcription factor

In our last example, we use the reduced Google matrix method in order to better understand the structure of regulations of known in advance direct targets of a selected transcription factor E2F1, a key transcription factor regulating cell cycle progression. For 76 such proteins found in SIGNOR pathway database, we find 103 direct interaction connecting these proteins into the connected component comprising 49 proteins. One additional direct transcriptional regulation was found between MYC and CBX5 proteins, but only in “cancer" K562 network.

The reduced Google matrix analysis revealed 84 indirect regulations for *G*_qr_ > 0.01 in the case of the “normal" network (Figure 8,A), all of which were also present in the “cancer" network. We did not find any additional indirect interactions from the analysis of the “cancer" network.

**Fig. 8.**
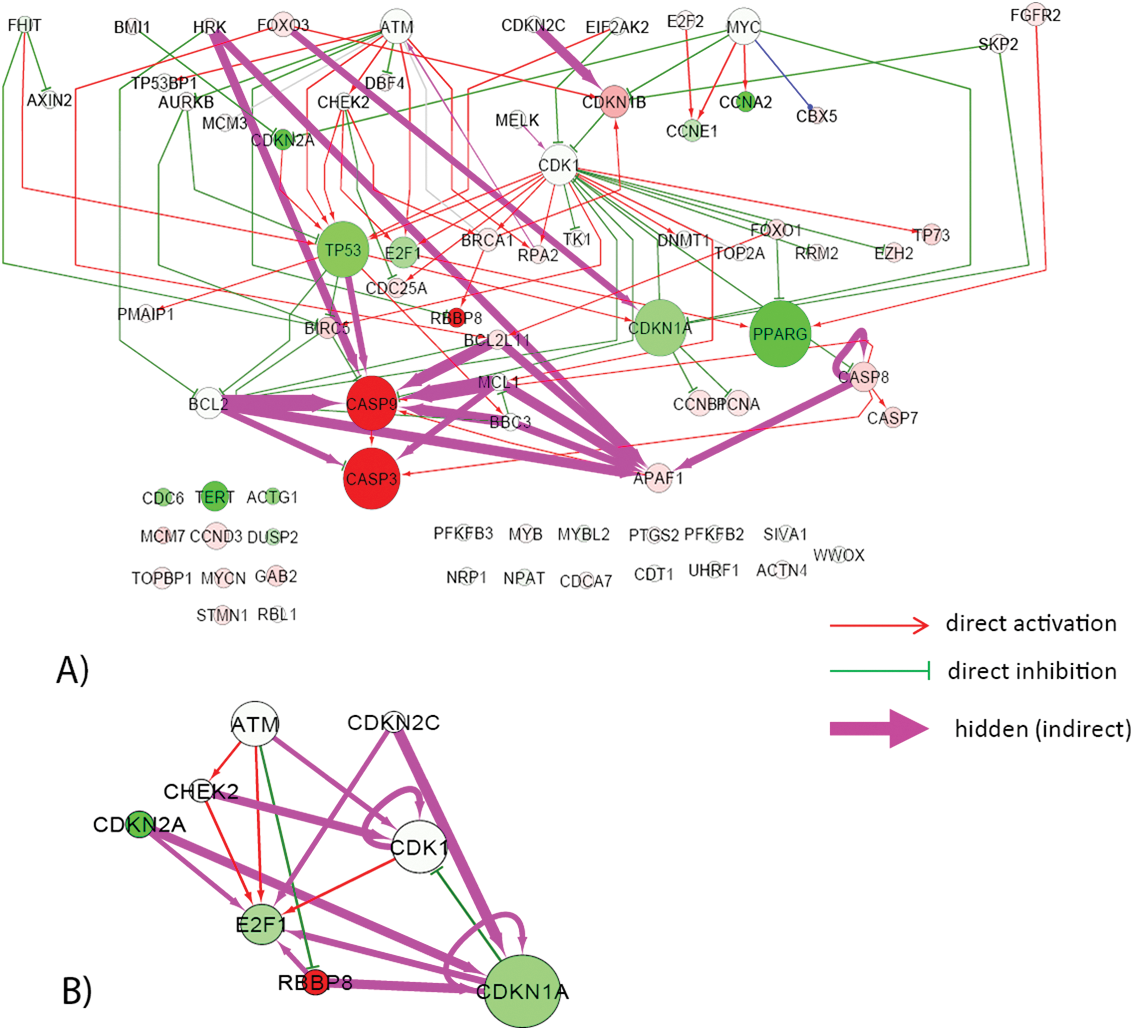
Network of targets of E2F1 transcription targets. The meaning of the node and edge colors is the same as in Figure 6. A) Complete network with a subset of 19 top scored indirect interactions is shown for *G*qr > 0.05. B) Network of E2F1 direct and indirect (*G*qr > 0.01) regulators of E2F1, showing multiple indirect feedback loops in the regulation of E2F1 itself.

The majority of strong indirect interactions pointed to the 3 key apoptosis proteins CASP9, CASP3 and APAF1 (apoptotic protease activating factor) whose local PageRanks decreased in the “cancer" network (which means they become more regulated). As in the AKT-mTOR example, we find indirect regulations between BCL2, BCL2L1 and CASP3. However, unlike AKT-mTOR example, we did not observe significant changes of local PageRanks of BCL2 and BCL2L1. Overall, the reduced Google matrix analysis underlines existence of hidden indirect apoptotic program regulated by E2F1 (which is a known fact [5]).

We also found that many weaker indirect interactions between the targets of E2F1 ends up on the E2F1 itself (Figure 8,B), also through a key G1/S cell cycle checkpoint protein CDKN1A (cyclin dependent kinase inhibitor 1A). Therefore, E2F1 itself can be regulated through a number of direct (3 in Figure 8,B) and even more indirect (4 in Figure 8,B) feedback regulations. This observation can provide hints on the principles of organization of the cell cycle transcriptional program.

## 3 Materials and Methods

### 3.1 Google matrix construction and properties

The Google matrix *G* of a directed network of *N* nodes is constructed from the adjacency matrix *A*_*ij*_ which has elements 1 if a protein (node) *j* points to a protein (node) *i* and zero otherwise. Then the matrix elements of *G* take the standard form [10,39]

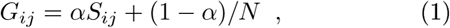

where *S* is the matrix of Markov transitions with elements 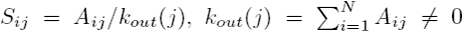 being the node *j* out-degree (number of outgoing links) and with *S*_*ij*_ = 1*/N* if *j* has no outgoing links (dangling node). Here 0 < *α* < 1 is the damping factor which for a random surfer determines the probability (1 − *α*) to jump to any node. The properties of spectrum and eigenstates of *G* have been discussed in detail for Wikipedia and other directed networks (see e.g. [20]).

The right eigenvectors *ψ*_*i*_(*j*) of *G* are determined by the equation:

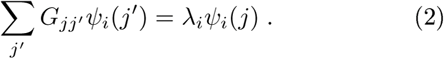

The PageRank eigenvector *P* (*j*) = *ψ*_*i*=0_(*j*) corresponds to the largest eigenvalue *λ*_*i*=0_ = 1 [10,39]. It has positive elements which give a probability to find a random surfer on a given node in the stationary long time limit of the Markov process. All nodes can be ordered by a monotonically decreasing probability *P*(*K*_*i*_) with the highest probability at *K* = 1. The index *K* is the PageRank index. Left eigenvectors are biorthogonal to right eigenvectors of different eigenvalues. The left eigenvector for *λ* = 1 has identical (unit) entries due to the column sum normalization of *G*. One can show that the damping factor *α* in (1) only affects the PageRank vector (or other eigenvectors for *λ* = 1 of *S* in case of a degeneracy) while other eigenvectors are independent of *α* due to their orthogonality to the left unit eigenvector for *λ* = 1 [39]. Thus all eigenvalues, except *λ* = 1, are multiplied by a factor *α* when replacing *S* by *G*. In the following we use the notations 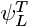 and *ψ*_*R*_ for left and right eigenvectors respectively (here *T* means vector or matrix transposition).

In many real networks the number of nonzero elements in a column of *S* is significantly smaller than the whole matrix size *N* that allows to find efficiently the PageRank vector by the PageRank algorithm of power iterations [39]. Also a certain number of largest eigenvalues (in modulus) and related eigenvectors can be efficiently computed by the Arnoldi algorithm (see [20] and Refs. therein).

In addition to the matrix *G* it is useful to introduce a Google matrix *G*^*∗*^ constructed from the adjacency matrix of the same network but with inverted direction of all links. The statistical properties of the eigenvector *P*^*∗*^ of *G*^*∗*^ with the largest eigenvalue *λ* = 1 have been studied first for the Linux Kernel network [15] showing that there are nontrivial correlations between *P* and *P*^*∗*^ vectors of the network. More detailed studied have been done for Wikipedia and other networks [20]. The vector *P*^*∗*^(*K*^*∗*^) is called the CheiRank vector and the index numbering nodes in order of monotonic decrease of probability *P*^*∗*^ is noted as CheiRank index *K*^*∗*^. Thus, nodes with many ingoing links have small value of *K* = 1, 2, 3, … and nodes with many outgoing links have *K*^*∗*^ = 1, 2, 3, … [39,20]. Examples of density distributions for Wikipedia editions and other directed networks are given in [20]. It is also useful to use 2DRank index *K*_2_ which represents a certain combination of *K*, *K*^*∗*^ indexes (*K*_2_ is the sequence of *K*, *K*^∗^ values appearing first on a sequence of squares which have left corner at *K* = *K*^*∗*^ = 1 with size increasing one by one up to maximal *N* value, see details in [20]).

At *α* < 1 only the PageRank vector have *λ* = 1 while all other eigenvectors of *G* have |*λ*| ≤ *α* [39,20]. For Wikipedia is was shown that the eigenvectors with a large modulus of *λ* select some specific communities of Wikipedia network [20]. However, a priory it is not possible to know what are the meanings of these communities. Thus other methods are required to determine effective interactions between *N*_*r*_ nodes of a specific subset (group) of the global network of a large size *N* ≫ *N*_*r*_.

In this work we apply the Google matrix analysis to the directed network of protein interactions from the cancer database SIGNOR [45], and two hybrid networks, constructed by merging SIGNOR to two transcriptional networks measured in normal blood cells and in cancer (leukemia).

The SIGNOR directed network contains *N* = 2432 proteins (nodes) with the total number of links *N*_*l*_ = 6569. In all our analysis we use the typical damping factor value *α* = 0.85 [39].

For the studied protein networks the dependencies of PageRank and CheiRank probabilities on rank indexes are shown in Figure 9. The decay of probabilities is approximately described by a power law *P* ∝ 1/*K*^*β*^; *P*^∗^ ∝ 1/*K*^∗*β*^ with the decay exponent *β* in a range 0.5 − 1. However, this is only an approximation for a whole curve. The distribution on nodes on the PageRank-CheiRank plane is shown in Figure 2. The spectra of *G* and *G*^*∗*^ are shown in Figure 10.

**Fig. 9.**
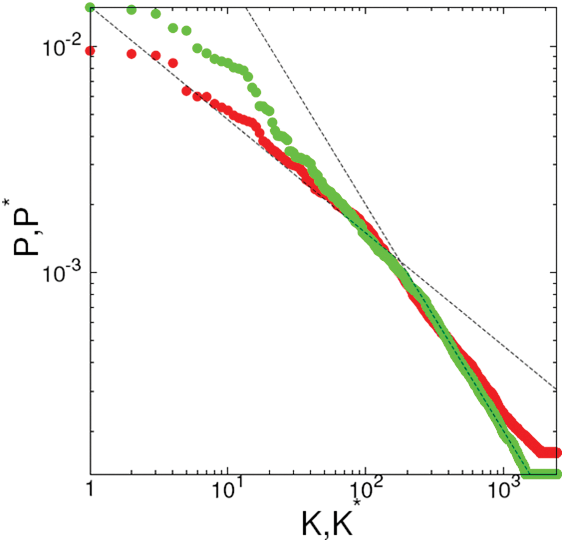
PageRank probability *P* (red points) and CheiRank probability *P*^*∗*^ (green points) as a function of *K* and *K*^*∗*^ indexes, calculated for SIGNOR molecular interaction database. Straight dashed lines are drawn to adapt an eye and show algebraic decay with exponents *−*0.5 and *−*1. Here *α* = 0.85.

**Fig. 10.**
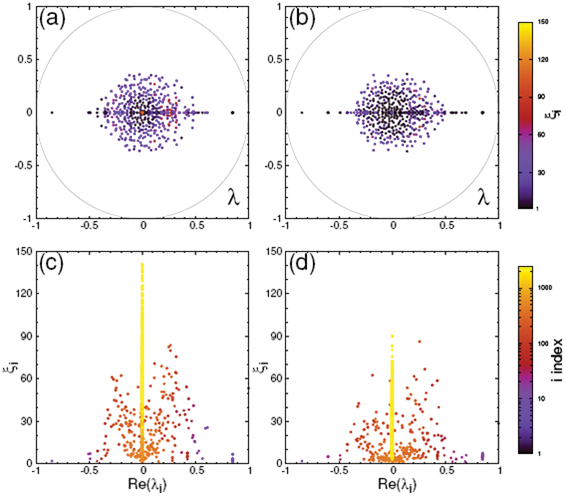
(a,b) Complex spectrum {*λ*_*i*_}_*i*=0,…,*N*−1_ of Google matrices *G* (a) and *G*^*∗*^ (b) at *α* = 0.85, computed for SIGNOR molecular interaction database. Colors give the inverse participation ratio *ξ*_*i*_ from *ξ*_*i*_ = 1 (black) to *ξ*_*i*_ = 150 (bright yellow). (c,d) Inverse participation ratio 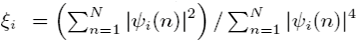 of the *i*th eigenvector as a function of Re (*λ*_*i*_) for Google matrices *G* (c) and *G*^*∗*^ (d). Colors give in logarithmic scale *i* indexes of eigenvalues ranked by decreasing modulus; from *i* = 1 (black) to *i* = *N* = 2432 (bright yellow). Quasi-uniform picture of the spectrum reflects inexistence of weak protein node communities in the SIGNOR network.

### 3.2 Reduced Google matrix

Recently, the method of reduced Google matrix has been proposed for analysis of effective interactions between nodes of a selected subset embedded into a large size network [21]. This approach uses parallels with the quantum scattering theory, developed for processes in nuclear and mesoscopic physics and quantum chaos.

It turns out that the Google matrix *G*_R_ matrix describing the interactions inside a group of proteins is composed of three matrix components which describe the direct interactions between group members, *G*_*rr*_, a projector part *G*_*pr*_ which is mainly imposed by the PageRank of selected proteins given by the global *G* matrix and a component *G*_qr_ from hidden interactions between proteins which appear due to indirect links via the global network. Thus the reduced matrix *G*_R_ = *G*_*rr*_+*G*_pr_+*G*_qr_ allows to obtain precise information about the group of proteins taking into account their environment given by the global network.

The concept of reduced Google matrix *G*_*R*_ was introduced in [21] on the basis of the following observation. At present directed networks of real systems can be very large (about 4.2 millions for the English Wikipedia edition in 2013 [20] or 3.5 billion web pages for a publicly accessible web crawl that was gathered by the Common Crawl Foundation in 2012 [40]). In certain cases one may be interested in the particular interactions among a small reduced subset of *N*_*r*_ nodes with *N*_*r*_ ≪ *N* instead of the interactions in the entire network. However, the interactions between these *N*_*r*_ nodes should be correctly determined taking into account that there are many indirect links between the *N*_*r*_ nodes via all other *N*_*s*_ = *N* − *N*_*r*_ nodes of the network. This leads to the problem of the reduced Google matrix *G*_*R*_ with *N*_*r*_ nodes which describes the interactions of a subset of *N*_*r*_ nodes.

In a certain sense we can trace parallels with the problem of quantum scattering appearing in nuclear and mesoscopic physics [47,4,30] and quantum chaotic scattering [26]. Indeed, in the scattering problem there are effective interactions between open channels to localized basis states in a well confined scattering domain where a particle can spend a certain time before its escape to open channels. Having this analogy in mind we construct the reduced Google matrix *G*_*R*_ which describes interactions between selected *N*_*r*_ nodes and satisfies the standard requirements of the Google matrix.

Let *G* be a typical Google matrix of Perron-Frobenius type for a network with *N* nodes such that *G*_*ij*_ ≥ 0 and the column sum normalization 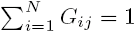 are verified. We consider a sub-network with *N*_*r*_ < *N* nodes, called “reduced network”. In this case we can write *G* in a block form :

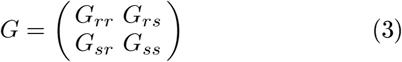

where the index “*r*” refers to the nodes of the reduced network and “*s*” to the other *N*_*s*_ = *N* − *N*_*r*_ nodes which form a complementary network which we will call “scattering network”.

We denote the PageRank vector of the full network as

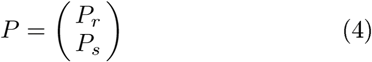

which satisfies the equation *G P* = *P* or in other words *P* is the right eigenvector of *G* for the unit eigenvalue. This eigenvalue equation reads in block notations:

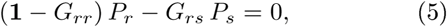

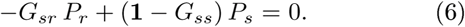

Here 1 is a unit diagonal matrix of corresponding size *N*_*r*_ or *N*_*s*_. Assuming that the matrix 1 − *G*_*ss*_ is not singular, i.e. all eigenvalues *G*_*ss*_ are strictly smaller than unity (in modulus), we obtain from (6) that

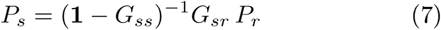

which gives together with (5):

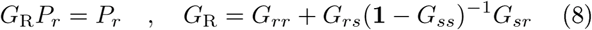

where the matrix *G*_R_ of size *N*_*r*_ × *N*_*r*_, defined for the reduced network, can be viewed as an effective reduced Google matrix. Here the contribution of *G*_*rr*_ accounts for direct links in the reduced network and the second term with the matrix inverse corresponds to all contributions of indirect links of arbitrary order. We note that in mesoscopic scattering problems one typically uses an expression of the scattering matrix which has a similar structure where the scattering channels correspond to the reduced network and the states inside the scattering domain to the scattering network [4].

The matrix elements of *G*_*R*_ are non-negative since the matrix inverse in (8) can be expanded as:

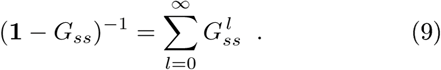

In (9) the integer *l* represents the order of indirect links, i. e. the number of indirect links which are used to connect indirectly two nodes of the reduced network. The matrix inverse corresponds to an exact resummation of all orders of indirect links. According to (9) the matrix (1−*G*_*ss*_)^−1^ and therefore also *G*_R_ have non-negative matrix elements. It can be shown that *G*_R_ also fulfills the condition of column sum normalization being unity [21].

The results obtained in [21,22] show that the reduced Google matrix can be presented as a sum of three components

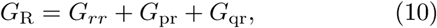

with the first component *G*_*rr*_ given by direct matrix elements of *G* among the selected *N*_*r*_ nodes, the second projector component *G*_pr_ is given by

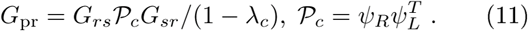

Here *λ*_*c*_ is the leading eigenvalue and by 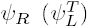 the corresponding right (left) eigenvector such that *G*_*ss*_ψ_*R*_ = *λ*_*c*_*ψ*_*R*_ (or 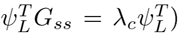). Both left and right eigenvectors as well as *λ*_*c*_ can be efficiently computed by the power iteration method in a similar way as the standard PageRank method. We note that one can easily show that *λ*_*c*_ must be real and that both left/right eigenvectors can be chosen with positive elements. Concerning the normalization for *ψ*_*R*_ we choose 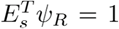 and for *ψ*_*L*_ we choose 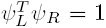 (the vector 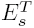 has all elements being unity). It is well known (and easy to show) that 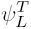 is orthogonal to all other right eigenvectors (and *ψ*_*R*_ is orthogonal to all other left eigenvectors) of *G*_*ss*_ with eigenvalues different from *λ*_*c*_. Here we introduce the operator 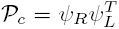 which is the projector onto the eigenspace of *λ*_*c*_ and we denote by 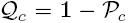 the complementary projector. One verifies directly that both projectors commute with the matrix *G*_*ss*_ and in particular 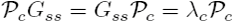.

We mention that this contribution is of the form 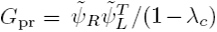 with 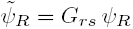 and 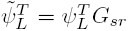 being two small vectors defined on the reduced space of dimension *N*_*r*_. Therefore *G*_pr_ is indeed a (small) matrix of rank one which is also confirmed by a numerical diagonalization of this matrix. The third component *G*_qr_ of indirect or hidden links is given by

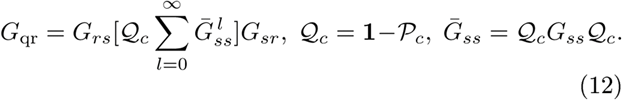

Even though the decomposition (10) is at first motivated by the numerical efficiency to evaluate the matrix inverse it is equally important concerning the interpretation of the different terms and especially the last contribution (12) which is typically rather small as compared to (11) plays in an important role as we will see below.

Concerning the numerical algorithm to evaluate all contributions in (10), we mention that we first determine by the power iteration method the leading left *ψ*_*L*_ and right eigenvector *ψ*_*R*_ of the matrix *G*_*ss*_ which also provides an accurate value of the corresponding eigenvalue *λ*_*c*_ or better of 1 − *λ*_*c*_ (by taking the norm of the projection of *Gψ*_*R*_ on the reduced space which is highly accurate even for *λ*_*c*_ close to 1). These two vectors provide directly *G*_pr_ by (11) and allow to numerically apply the projector 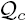 to an arbitrary vector (with ∼ *N* operations). The most expensive part is the evaluation of the last contribution according to (12). For this we apply successively *G*_*ss*_ and 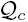 to an arbitrary column of *G*_*sr*_ which can be done by a sparse matrix vector multiplication or the efficient application of the projector. We compute simultaneously the series in (12) which converges rather quickly after about 200 terms since the contribution of the leading eigenvalue (of *G*_*ss*_) has been taken out and the eigenvalues of 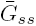 are roughly below the damping factor *α* = 0.85. In the end the resulting vector is multiplied with the matrix *G*_*rs*_ which provides one column of *G*_qr_. This procedure has to be repeated for each of the *N*_*r*_ columns but the number *N*_*r*_ is typically rather modest. We also note that the results obtained in [22] show that an approximate relation holds: 1 − *λ*_*c*_ ≈ *Σ*_*P*_ = ‖*P*_*r*_‖_1_ where *Σ*_*P*_ is the PageRank probability of the global network concentrated on the subset of *N*_*r*_ selected nodes.

The results obtained here and in [22] for the Wikipedia network show that the contribution of *G*_pr_ is dominant in *G*_*R*_ but it is also kind of trivial with nearly identical columns. Therefore the two small contributions of *G*_*rr*_ and *G*_qr_ are indeed very important for the interpretation even though they only contribute weakly to the overall column sum normalization.

The meaning of *G*_*rr*_ is rather clear since is gives direct links between the selected nodes. In contrast, the meaning of *G*_qr_ is significantly more interesting since it generates indirect links between the *N*_*r*_ nodes due to their interactions with the global network environment. We note that *G*_qr_ is composed of two parts *G*_qr_ = *G*_qrd_+*G*_qrnd_ where the first diagonal term *G*_qrd_ represents a probability to stay on the same node during multiple iterations of 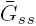 in (12) while the second nondiagonal term *G*_qrnd_ represents indirect (hidden) links between the *N*_*r*_ nodes appearing due via the global network. We note that in principle certain matrix elements of *G*_qr_ can be negative, which is possible due to negative terms in 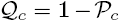 appearing in (12). However, for all subsets considered in this work the total weight of negative elements was negligibly small (at most some 10^−3^) of the total weight 1 for *G*_*R*_).

It is convenient to characterize the strength of 3 components in (10) by their respective weights *W*_*rr*_, *W*_pr_, *W*_qr_ given respectively by the sum of all matrix elements of *G*_*rr*_, *G*_pr_, *G*_qr_ divided by *N*_*r*_. By definition we have *W*_*rr*_ + *W*_pr_ + *W*_qr_ = 1. All numerical data of the reduced Google matrix of groups of proteins considered here are publicly available at the web site [52].

### 3.3 Global network reconstruction for GM12878 and K562 cell lines

The transcriptional networks for normal GM12878 and cancer K562 cell lines were obtained from the web-site http://encodenets.gersteinlab.org/ (files enets7.K562_proximal_filtered_network.txt, enets8.GM_proximal_filtered_network.txt) accompanying the original publication [27]. SIGNOR network for H.Sapiens was downloaded from the SIGNOR web-site http://signor.uniroma2.it/downloads.php. Both transcriptional and SIGNOR networks were represented as simple interaction format (SIF) files and merged by simple concatenation. They were further processed in in Cytoscape [16] with use of BiNoM plugin [50,9] for finding shortest and second shortest paths, and copy-paste operations.

### 3.4 Deinitions of functionally related groups of proteins

The composition of AKT-mTOR pathway and the set of direct transcriptional targets of E2F1 protein were downloaded from the Atlas of Cancer Signaling Network (ACSN) database [37], by using GMT files of version 1.1 available from the ACSN web-site http://acsn.curie.fr (sets E2F1-TARGETS and AKT-mTOR gene sets).

The group of proteins related to proliferation was determined as a set of 50 gene names top-contributing to the transcriptomic signature associated to the cell cycle through a large-scale pan-cancer analysis of transcriptomic data [8], using the lists provided in the Supplementary information [52].

## 4 Discussion

The results of application of high-throughput technologies in modern molecular biology are more and more frequently presented in the form of complex networks, representing measured causal relations between biological molecules. For example, systematic application of Chip-Seq technology for a significant number of transcription factors can result in the global cell line-specific reconstruction of the transcriptional network [27]. Despite many methods aimed at the analysis of complex networks, there is still a need for mathematically rigorous and computationally efficient methods able to quantify complex non-local network topologies, especially in the case of directed networks.

In this work we show that the global Google matrix and the reduced Google matrix approaches represent useful tools for the analysis of directed interaction networks in biology.

We show that the global analysis of a directed biological network using Google matrix and by computing node PageRanks and CheiRanks and their relative changes in cancer allows obtaining insights about specific and precise aspects of how the biological network topology evolve in different biological contexts.

The reduced Google matrix approach is a novel method allowing quantifying indirect (hidden) connections between members of a specified subset of network nodes. These connections represent paths of oriented graph edges through the global network and involving nodes outside the specified set. This approach is applied to the global network of directed protein-protein interactions, with a focus on some groups of proteins corresponding to a well-defined biological function (cell survival signaling, cell proliferation), obtained by different methods (prior knowledge-based or by data-driven approaches). We show that application of the reduced Google matrix approach leads to inferring a meaningful set of indirect interactions highlighting existence of specific biological programs not reflected in the structure of direct relations between the members of a protein set. We also show that the structure of such hidden relations can be modified from one condition to another, reflecting some global changes in the wiring of, for example, global transcriptional networks during cancer or differentiation.

There are multiple possible ways to exploit the methods suggested in this study. One of the promising application is in mathematical modeling of biological processes, i.e. mathematical modeling of molecular pathways [12,29,17,44]. Construction of a mathematical model of a pathway usually starts with defining the restrictive set of biological molecules or processes most closely related to the studied phenomenon (i.e., regulation of programmed cell death). In the current methodologies, the number of such model elements (proteins) can not be very large. Therefore, there is always a danger of neglecting important indirect causal relations between the elements via regulations passing through the global network in which a given pathway is embedded. The reduced Google matrix method allows systematically inferring indirect regulations, in a context-specific manner which allows to use in this analysis the results of high-throughput biotechnologies, as it is demonstrated in the current study.

Note that indirect regulations can involve too many proteins in order to characterize them by *ad hoc* methods such as counting the number of paths connecting two proteins. The suggested method takes into account the directed network in its whole complexity without naive simplifying assumptions. Moreover, the method is computationally efficient for the typical sizes of the biological networks involving tens of thousands of nodes and hundreds of thousands of interactions.

Google matrix, or Googlomics, methodology can be used in other types of directed networks appearing in biology such as state transition graphs resulting from the analysis of Boolean models of pathways [1].

Overall, the developed methodology allows combining global structural analysis of large biological networks characterized by context-specific and dynamical re-wiring together with the focused analysis of specified biological processes, without neglecting the role of the global context.

## 5 Acknowledgements

This research is supported in part by the MASTODONS-2016 CNRS project APLIGOOGLE (see http://www.quantware.ups-tlse.fr/APLIGOOGLE/).

## References

1. W. Abou-Jaoudé, P. Traynard, P.T. Monteiro, J. Saez-Rodriguez, T. Helikar, D. Thieffry, C. Chaouiya. Logical Modeling and Dynamical Analysis of Cellular Networks. Front Genet. 7:94 (2016)

2. L. Albergante, J. Blow, T.J. Newman Buffered Qualitative Stability explains the robustness and evolvability of transcriptional networks Elife 3:e02863 (2014)

3. E. Barillot, L. Calzone, P.Hupe, J.-P. Vert and A. Zinovyev. Computational systems biology of cancer, CRC Press, 2012.

4. C.W.J. Beenakker, Random-matrix theory of quantum transport, Rev. Mod. Phys. 69, 731 (1997)

5. L.A. Bell, J. O’Prey, K.M. Ryan. DNA-binding independent cell death from a minimal proapoptotic region of E2F-1. Oncogene 25(41):5656–63 (2006).

6. J.D. West, T.C. Bergstrom and C.T. Bergstrom, Big Macs and Eigenfactor Scores: Don’t Let Correlation Coefficients Fool You J. American Society Iform. Sci. Technology 61:1800–1807 (2010); http://eigenfactor.org

7. M. Rosvall, C.T. Bergstrom. Maps of random walks on complex networks reveal community structure. Proceedings of the National Academy of Sciences 105:(4), 1118–1123 (2008).

8. A. Biton, I. Bernard-Pierrot, Y. Lou, C. Krucker, E. Chapeaublanc, C. Rubio-Pérez, N. López-Bigas, A. Kamoun, Y. Neuzillet, P. Gestraud, L. Grieco, S. Rebouissou, A. de Reyniès, S. Benhamou, T. Lebret, J. Southgate, E. Barillot, Y. Allory, A. Zinovyev, F. Radvanyi. Independent component analysis uncovers the landscape of the bladder tumor transcriptome and reveals insights into luminal and basal subtypes. Cell Rep. 9(4):1235–1245 (2014)

9. E. Bonnet, L. Calzone, D. Rovera, G. Stoll, E. Barillot and A. Zinovyev BiNoM 2.0, a Cytoscape plugin for accessing and analyzing pathways using standard systems biology formats BMC Systems Biology 7:18, (2013)

10. S. Brin and L. Page, The anatomy of a large-scale hypertextual Web search engine, Computer Networks and ISDN Systems 30, 107 (1998)

11. L. Calzone, A. Gelay, A. Zinovyev, F. Radvanyi, E. Barillot. A comprehensive modular map of molecular interactions in RB/E2F pathway. Mol Syst Biol. 4:173 (2008)

12. L. Calzone, L. Tournier, S. Fourquet, D. Thieffry, B. Zhivotovsky, E. Barillot, A. Zinovyev. Mathematical modelling of cell-fate decision in response to death receptor engagement. PLoS Comput Biol 5;6(3):e1000702, (2010)

13. J. Chen, E.E. Bardes, B.J. Aronow, A.G. Jegga. ToppGene Suite for gene list enrichment analysis and candidate gene prioritization. Nucleic Acids Research 37(suppl 2):W305–W311 (2009)

14. L. Chindelevitch, D. Ziemek, A. Enayetallah, R. Randhawa, B. Sidders, C. Brockel and E.S. Huang. Causal reasoning on biological networks: interpreting transcriptional changes Bioinformatics 288, 1114–1121 (2012)

15. A.D. Chepelianskii, Towards physical laws for software architecture, arXiv:1003.5455 [cs.SE] (2010)

16. M.S. Cline, M. Smoot, E. Cerami, A. Kuchinsky, N. Landys, C. Workman, R. Christmas, I. Avila-Campilo, M. Creech, B. Gross, K. Hanspers, R. Isserlin, R. Kelley, S. Killcoyne, S. Lotia, S. Maere, J. Morris, K. Ono, V. Pavlovic, A.R. Pico, A. Vailaya, P.L. Wang, A. Adler, B.R. Conklin, L. Hood, M. Kuiper, C. Sander, I. Schmulevich, B. Schwikowski, G.J. Warner, T. Ideker, G.D. Bader. Integration of biological networks and gene expression data using Cytoscape. Nat Protoc. 2(10):2366–2382 (2007)

17. P.A.D. Cohen, L. Martignetti, S. Robine, E. Barillot, A. Zinovyev, L. Calzone. Mathematical Modelling of Molecular Pathways Enabling Tumour Cell Invasion and Migration. PLoS Computational Biology 11:e1004571, (2015)

18. P. Csermely, T. Korcsmáros, H.J. Kiss, G. London, R. Nussinov. Structure and dynamics of molecular networks: a novel paradigm of drug discovery: a comprehensive review. Pharmacol Ther. 138(3):333–408 (2013)

19. Q. Cui, Y. Ma, M. Jaramillo, H. Bari, A. Awan, S. Yang, S. Zhang, L. Liu, M. Lu, M. O’Connor-McCourt, E.O. Purisima, E. Wang. A map of human cancer signaling. Mol Syst Biol. 3:152 (2007)

20. L. Ermann, K.M. Frahm and D.L. Shepelyansky, Google matrix analysis of directed networks, Rev. Mod. Phys. 87, 1261 (2015)

21. K.M. Frahm and D.L. Shepelyansky, Reduced Google matrix, arXiv:1602.02394[physics.soc] (2016)

22. K.M. Frahm, K. Jaffrès-Runser and D.L. Shepelyansky, Wikipedia mining of hidden links between political leaders, Eur. Phys. J. B 89, 269 (2016)

23. K.M. Frahm, B. Georgeot and D.L. Shepelyansky, Universal emergence of PageRank, J. Phys. A: Math. Theor. 44, 465101 (2011)

24. D. Fazekas, M. Koltai, D. Türei, D. Módos, M. Pálfy, Z. Dúl, L. Zsákai, M. Szalay-Bek, K. Lenti, I.J. Farkas, T. Vellai, P. Csermely, T. Korcsmáros. SignaLink 2 – a signaling pathway resource with multi-layered regulatory networks. BMC Systems Biology 7(1):7 (2013)

25. M. Garber, N. Yosef, A. Goren, R. Raychowdhury, A. Thielke, M. Guttman, J. Robinson, B. Minie, N. Chevrier, Z. Itzhaki, R. Blecher-Gonen, C. Bornstein, D. Amann-Zalcenstein, A. Weiner, D. Friedrich, J. Meldrim, O. Ram, C. Cheng, A. Gnirke, S. Fisher, N. Friedman, B. Wong, B.E. Bernstein, C. Nusbaum, N. Hacohen, A. Regev, I. Amit. A high-throughput chromatin immunoprecipitation approach reveals principles of dynamic gene regulation in mammals. Mol Cell. 47(5):810–22 (2012).

26. P. Gaspard, Quantum chaotic scattering, Scholarpedia 9(6), 9806 (2014)

27. MB Gerstein, A Kundaje, M Hariharan et al. Architecture of the human regulatory network derived from ENCODE data Nature 489, 91–100 (2012).

28. G. Giordano, C. C. Samaniego, E. Franco and· F. Blanchini. Computing the structural influence matrix for biological systems J. Math. Biol. 72(7),1927–1958 (2015)

29. L. Grieco, L. Calzone, I. Bernard-Pierrot, F. Radvanyi, B. Kahn-Perles, D. Thieffry. Integrative modelling of the influence of MAPK network on cancer cell fate decision. PLoS Comput. Biol. 9, e1003286 (2013)

30. T. Guhr, A. Müller-Groeling and H.A. Weidenmüller, Random Matrix Theories in Quantum Physics: Common Concepts, Phys. Rep. 299, 189 (1998)

31. M. Hofree, J.P. Shen, H. Carter, A. Gross, T. Ideker, Network-based stratification of tumor mutations Nat Methods 1011, 1108–1115 (2013)

32. H. Jeong, S.P. Mason, A.-L. Barabási and Z.N. Oltvai. Lethality and centrality in protein networks. Nature, 411:41–42 (2001).

33. S. Köhler, S. Bauer, D. Horn, P.N. Robinson. Walking the interactome for prioritization of candidate disease genes. Am. J. Hum. Genet. 82, 949—958, (2008)

34. K. Komurov, M.A. White, P.T. Ram Use of Data-Biased Random Walks on Graphs for the Retrieval of Context-Specific Networks from Genomic Data PLoS Computational Biology 6(8),e1000889 (2010)

35. J. Koschmann, A. Bhar, P. Stegmaier, A.E. Kel and E. Wingender. “Upstream Analysis”: An Integrated Promoter-Pathway Analysis Approach to Causal Interpretation of Microarray Data Microarrays 4, 270–286 (2015)

36. I. Kuperstein, L. Grieco, D.P. Cohen, D. Thieffry, A. Zinovyev, E. Barillot. The shortest path is not the one you know: application of biological network resources in precision oncology research. Mutagenesis 30(2), 191–204 (2015)

37. I. Kuperstein, E. Bonnet, H.A. Nguyen, D. Cohen, E. Viara, L. Grieco, S. Fourquet, L. Calzone, C. Russo, M. Kondratova, M. Dutreix, E. Barillot, A. Zinovyev. Atlas of Cancer Signalling Network: a systems biology resource for integrative analysis of cancer data with Google Maps. Oncogenesis 4:e160, (2015)

38. I. Lee, U.M. Blom, P.I. Wang, J.E. Shim, and E.M. Marcotte. Prioritizing candidate disease genes by network-based boosting of genomewide association data. Genome Res. 21, 1109—1121 (2011)

39. A.M. Langville and C.D. Meyer, Google’s PageRank and beyond: the science of search engine rankings, Princeton University Press, Princeton (2006)

40. R. Meusel, S. Vigna, O. Lehmberg and C. Bizer, The graph structure in the web - analyzed on different aggregation levels, J. Web Sci. 1, 33 (2015)

41. R. Pique-Regi, J.F. Degner, A.A. Pai, D.J. Gaffney, Y. Gilad, J.K. Pritchard. Accurate inference of transcription factor binding from DNA sequence and chromatin accessibility data. Genome Res. 21(3):447–55 (2011)

42. M.A. Pujana, J.D. Han, L.M. Starita, et al. (2007) Network modeling links breast cancer susceptibility and centrosome dysfunction. Nat. Genet. 39, 1338—1349 (2007)

43. S. Redner (2005) Citation statistics from 110 years of Physical Review. Phys. Today 58(6), 49—54 (2005)

44. E. Remy, S. Rebouissou, C. Chaouiya, A. Zinovyev, F. Radvanyi, L. Calzone. A Modeling Approach to Explain Mutually Exclusive and Co-Occurring Genetic Alterations in Bladder Tumorigenesis. Cancer Res. 75(19):4042–4052, (2015)

45. L. Perfetto, L. Briganti, A. Calderone et al. SIGNOR: a database of causal relationships between biological entities. Nucleic Acids Res. 44(D1):D548–54 (2016)

46. F. Rapaport, A. Zinovyev, M. Dutreix, E. Barillot, J.P. Vert. Classification of microarray data using gene networks, BMC Bioinformatics 8:35 (2007)

47. V.V. Sokolov and V.G. Zelevinsky, Dynamics and statistics of unstable quantum states, Nucl. Phys. A 504, 562 (1989)

48. O. Vanunu, O. Magger, E. Ruppin, T. Shlomi, R. Sharan. Associating genes and protein complexes with disease via network propagation. PLoS Comput. Biol. 6, e1000641 (2010)

49. C. Winter, G. Kristiansen, S. Kersting, J. Roy, D. Aust, T. Knösel, P. Rümmele, B. Jahnke, V. Hentrich, F. Rückert, M. Niedergethmann, W. Weichert, M. Bahra, H.J. Schlitt, U. Settmacher, H. Friess, M. Büchler, H.D. Saeger, M. Schroeder, C. Pilarsky, R. Grützmann. Google goes cancer: improving outcome prediction for cancer patients by network-based ranking of marker genes. PLoS Comput Biol. 8(5):e1002511. (2012)

50. A. Zinovyev, E. Viara, L. Calzone, E. Barillot. BiNoM: a Cytoscape plugin for manipulating and analyzing biological networks. Bioinformatics 24(6):876–877 (2008)

51. A. Zinovyev, U. Kairov, T. Karpenyuk, E. Ramanculov. Blind source separation methods for deconvolution of complex signals in cancer biology. Biochem Biophys Res Commun. 430(3):1182–1187 (2013)

52. http://www.quantware.ups-tlse.fr/QWLIB/googlomics/

